# Whole-community translocation shows high survival potential of pollution-sensitive benthic macroinvertebrate taxa in restored streams

**DOI:** 10.64898/2026.09.22.753605

**Authors:** R. Fagbohun Ibrahim, N. Sweetman Jon, H. Hilderbrand Robert, C. Allen Daniel

## Abstract

**Introduction:** Despite decades of investment, stream restoration often fails to recover diverse benthic macroinvertebrate communities, as restored reaches tend to remain dominated by pollution-tolerant taxa. This shortfall in recovery is commonly attributed to either poor habitat quality or dispersal limitation of aquatic macroinvertebrate taxa, yet these hypotheses are rarely tested directly.

**Objectives:** We conducted a manipulative field experiment in the eastern Piedmont region (Maryland, USA) within the Chesapeake Bay Watershed to evaluate whether restored streams can support pollution-sensitive taxa when dispersal barriers are removed.

**Methods:** We translocated whole macroinvertebrate communities from three nearby reference streams into three restored streams using standardized habitat substrates and quantified macroinvertebrate survival after 28 days. We also assessed how effective leaf vs rock substrates were for accumulating diverse and abundant macroinvertebrate communities for translocation.

**Results:** Across all sites, 77% of translocated individuals and 16 of 22 taxa sensitive and moderately sensitive to pollution persisted under restored stream conditions, demonstrating that restored reaches can support short-term survival of pollution-sensitive taxa that typically fail to recover. Substrate comparisons revealed that leaf packs accumulated higher macroinvertebrate abundances, whereas both substrates accumulated macroinvertebrate communities with similar taxa richness, EPT richness, and Shannon Diversity.

**Conclusions:** This work suggests that dispersal limitation, rather than unsuitable habitat, may be an underappreciated constraint limiting macroinvertebrate recovery in restored streams. These findings challenge the assumption that habitat restoration alone is sufficient for biological recovery and highlight the potential for whole-community translocations to accelerate recovery and ecological uplift as part of stream restoration practices.

**Implication for Practice:** Our study highlights the need to consider habitat connectivity and dispersal pathways during stream restoration designs. Restoring streams to meet habitat requirements of pollution-sensitive macroinvertebrate taxa may not be sufficient for their populations to recover if these taxa are unable to reach the restored stream. In scenarios where restoring connectivity to source populations is infeasible, whole community translocations may offer a novel restoration tool to overcome dispersal barriers and accelerate biological recovery in restored streams.

## Introduction

The diversity and taxonomic composition of benthic macroinvertebrate communities have long been used as an indicator of stream ecosystem health, as macroinvertebrate taxa respond to stream pollution in different ways, creating a gradient of pollution tolerance across taxa (Herman & Nejadhashemi 2015). In healthy streams with minimal pollution, pollution-sensitive and pollution-tolerant taxa co-occur, creating a functionally diverse and productive ecosystem capable of supporting a wide range of ecological processes (Houghton 2021). As streams become degraded, pollution-sensitive taxa disappear, leaving invertebrate communities dominated by pollution-tolerant taxa. This shift reduces functional diversity and ecological complexity, diminishing the capacity of streams to provide essential ecosystem services (Akamagwuna et al. 2021; Paz et al. 2022).

To reverse the widespread degradation of stream ecosystems and recover lost ecosystem services, stream restoration practices have become a widely adopted solution (Lake et al. 2007; Palmer et al. 2010). Typically, stream restoration focuses on using engineering methods to design and construct stable stream channels that structurally resemble natural streams (Johnson et al. 2020). The underlying assumption is that repairing the physical structure of the stream ecosystem will lead to the recovery of the biotic component, including benthic macroinvertebrates. This assumption has been referred to as the Field of Dreams Hypothesis (Palmer et al. 1997; Hilderbrand et al. 2005; Lake et al. 2007): because “*if you build it, they will come.”* Yet, benthic macroinvertebrate communities often remain depauperate following stream restorations, dominated by a few pollution-tolerant taxa, devoid of pollution-sensitive taxa, and generally similar to those of polluted streams, even several years following restoration (Louhi et al. 2011; Haase et al. 2013a).

Several explanations have been proposed to explain the poor recovery of benthic macroinvertebrate biodiversity in restored streams despite improvements in water quality and physical habitat. One explanation suggests that common restoration practices may not improve water quality enough to support the requirements of desired macroinvertebrate taxa (Haase et al. 2013). This explanation is grounded in a fundamental concept of community assembly known as environmental filtering or community sorting, and it describes how environmental factors strongly influence community structure (Cantera et al. 2023). Although there is a consensus in the literature that stream restoration improves habitat quality by reducing sedimentation and nutrient load (Thompson et al. 2018a), improving water chemistry and habitat heterogeneity (Frainer et al. 2018), and diversifying flow patterns and substrate composition (Stoll et al. 2016), there is little evidence that these new environmental conditions can support desired sensitive macroinvertebrate taxa.

In addition to insufficient improvements in habitat quality, another explanation is dispersal limitation, such that sensitive species cannot disperse well enough to restored streams to reestablish viable populations from source populations (Sundermann et al. 2011). The importance of dispersal and habitat connectivity is well known in ecological community assembly and is emphasized in metacommunity theory as proposed by Leibold et al. (2004). If restoration measures effectively alleviate environmental stressors and restored streams become hospitable to sensitive macroinvertebrate taxa, the ability of these taxa to disperse to restored streams plays a key role in their colonization and persistence. Unfortunately, most pollution-sensitive macroinvertebrate taxa, especially the Ephemeroptera-Plecoptera-Trichoptera (EPT) taxa, are weak fliers as adults and passive dispersers as larvae that rely on downstream current for movement from upstream populations not existing above restored segments (Petersen et al. 2004).

Recognizing the importance of habitat quality and dispersal, several studies have explored methods for including these regional-scale processes in stream restoration. Dispersal limitation may be overcome by translocating macroinvertebrate communities from unpolluted streams to restored streams to facilitate biodiversity recovery (Haase & Pilotto 2019; Dumeier et al. 2020; Macneale 2020). However, the success of such efforts would require sensitive macroinvertebrate taxa to be able to survive in the restored streams, which can be an open question. Here, we conducted a manipulative field experiment to investigate the survivability of pollution-sensitive macroinvertebrate taxa in three restored streams across the piedmont region of Maryland, USA. We address two specific questions: 1) Can pollution-sensitive benthic macroinvertebrates survive in restored streams upon translocation? 2) How can morphologically and functionally diverse and abundant macroinvertebrate communities be effectively translocated from source populations to restored streams?

## Methods

### Study Area

The study employs a manipulative field experiment to investigate the survival of sensitive benthic macroinvertebrate taxa in restored streams. We selected three restored streams across three different watersheds in the Eastern Piedmont physiographic region of Maryland. The restored streams receiving translocated macroinvertebrates are hereafter referred to as “recipient streams.” Recipient streams were selected because they were restored within the last 15 years while macroinvertebrate diversity and abundance remain low in recent bioassessments. Each of the recipient sites was paired with a suitable “donor stream”, which is a nearby reference stream with a Benthic Index of Biotic Integrity (BIBI) score > 4 that indicates a diverse and healthy macroinvertebrate assemblage (Figure 1). Donor and recipient streams were identified and selected using data in the Maryland Biological Stream Survey (MBSS) database (Hodgson et al. 2024). We paired recipient streams with donor streams that were within the same watershed or adjacent watersheds to minimize dissimilarities in ecological characteristics and natural macroinvertebrate distribution between donor and recipient sites, and also to prevent translocation of invasive species or pathogens across watersheds (Clinton et al. 2022).

**Figure 1:**
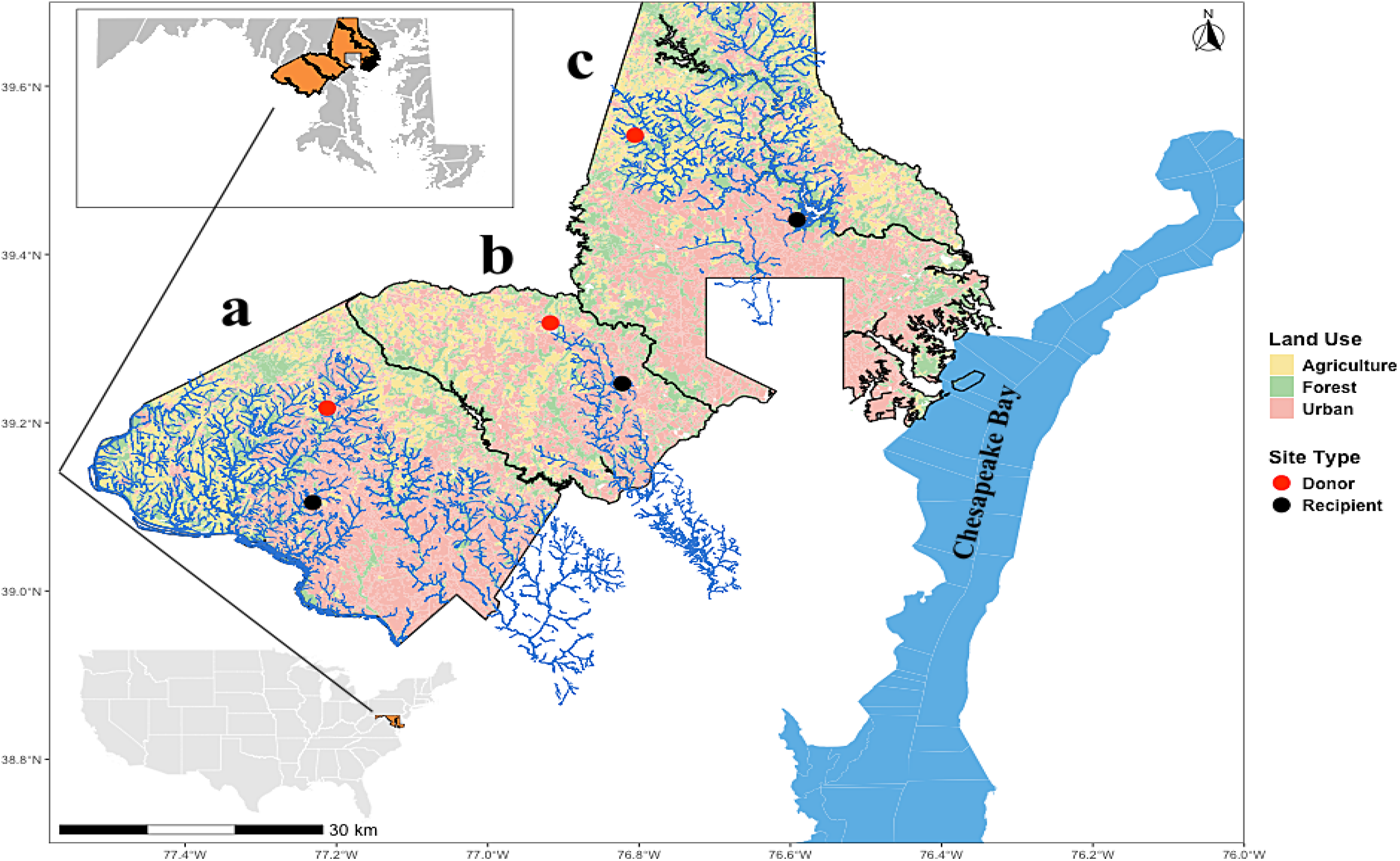
Map of study area showing the locations of the recipient and donor sites across three watersheds (a) Potomac River, (b) Little Patuxent, and (c) Loch Raven in the eastern Piedmont physiographic region of Maryland, USA.

### Macroinvertebrate translocations

At each donor site, we passively collected macroinvertebrate samples by deploying “habitat tubes”, open cylinders filled with natural substrate along a 100 m section of the stream to allow for passive natural colonization (Dumeier et al. 2018). Habitat tubes were made from Polyvinyl chloride (PVC) pipe sections, which provided support for an exterior mesh with 1.27 cm openings. Each tube has a length-width dimension of 76.2 by 22.8 cm, with an approximate volume of 31,112.9 cm^3^. Each tube was filled with either commercially sourced river rocks (size ranging between 7.62–12.7 cm) or abscised oak leaves sourced from residential neighborhoods in State College, PA (Figure S1).

In each donor stream, we deployed 20 habitat tubes, 10 filled with rocks and 10 filled with oak leaves. Habitat tubes were secured with zip ties to a steel rebar stake hammered into the stream bed where the water was deep enough to fully submerge the tubes. At each stake, we placed one leaf tube and one rock tube to ensure that both substrates were exposed to similar microhabitats. Habitat tubes were spatially distributed across different microhabitats, including riffles, pools, runs, and edges, and were additionally secured by a steel cable with a diameter of 3.175 mm attached to a tree on the streambank. Habitat tubes were deployed once in the fall and again in the spring to accumulate a wide range of taxa active in both seasons. In both seasons, habitat tubes were deployed for four weeks to allow enough time for macroinvertebrate colonization (Dumeier et al. 2018).

After the colonization phase, we randomly selected half of the habitat tubes from each donor stream and harvested them to characterize the macroinvertebrate communities before translocation. These harvested tubes served as pre-translocation controls, under the assumption that the community structure, diversity, and abundance of macroinvertebrates in the harvested tubes represent those in the tubes that were subsequently translocated. The other half were translocated to the recipient restored streams. When harvested, tubes were cut open, and their contents (substrate + macroinvertebrates) were emptied into an 11.36 L bucket half-filled with stream water. Large substrate particles were thoroughly washed to remove attached macroinvertebrates and were removed from the slurry. Once the bucket contained a slurry of small substrate particles, macroinvertebrates, and water, the slurry was passed through a 500 µm sieve. Sieve contents were then placed in a labelled sample jar filled with 95% ethanol and transported to the lab for sorting and identification. The tubes selected for translocation were removed from the stream and wrapped in a fine mesh (mesh size 1.016 mm) to prevent loss of macroinvertebrate from the habitat tubes during transport. The enclosed tubes were placed in coolers filled with stream water and fitted with portable aerators to maintain cool and oxygenated conditions during transport to recipient sites.

At the recipient sites, the tubes remained covered with fine mesh (1.016 mm) to prevent immigration and emigration of macroinvertebrates, while still allowing some natural water flow and particulate transport into the tubes. The tubes were placed along a 100 m stretch of the recipient stream in selected locations where the water was deep enough to submerge the cages, but with sufficient streamflow to mirror the habitat conditions selected at the donor streams. The tubes were secured as earlier described for the donor sites. After four weeks, the habitat tubes were removed from the recipient sites and harvested as described above to assess the survival of macroinvertebrate taxa following translocation. Finally, to assess the effect of seasonality on macroinvertebrate colonization and survival, we conducted translocations once in the fall of 2024 and once in the spring of 2025.

### Macroinvertebrate processing

In the lab, we removed alcohol from the macroinvertebrate jars by running the samples through a 500 mm sieve. Sieve contents were then emptied into a white tray and submerged in water for picking. Macroinvertebrates were carefully picked out of the tray and counted into a vial filled with 95% ethanol. For samples that exceeded 600 individuals, we subsampled using a Folsom plankton splitter (Sell & Evans 1982) that divides a sample into two approximately equal parts. We subsampled until the number of individuals fell between 300 and 500, following Chen et al. (2015). Identifications were made to genus level except for Chironomidae, which were identified to the family level.

### Statistical analysis - community composition

We used a multivariate ordination analysis to compare macroinvertebrate community composition in habitat tubes harvested at the end of the colonization period at the donor sites and those harvested at the end of the survival period at the recipient sites, which hereafter we refer to as “pre-translocation” and “post-translocation”, respectively. Before conducting the multivariate ordination analysis, we separated the dataset into fall and spring samples and analyzed each independently to avoid the pronounced seasonal difference in community composition overwhelming the treatment-related effects.

We calculated Bray-Curtis distances, then used Non-metric Multidimensional Scaling (NMDS) to ordinate the communities. Before calculating the Bray-Curtis distance and running the ordination, we square-root transformed raw abundance data to reduce the dominance of highly abundant taxa such as *Allocapnia*, Chironomidae, and *Stenonema.* We also removed rare taxa, defined as those not found in at least three pre-translocation tubes. To test for a significant difference in community centroids between pre-translocation and post-translocation macroinvertebrate communities, we used Permutational Multivariate Analysis of Variance (PERMANOVA, (Anderson & Walsh 2013). After running the PERMANOVA, we tested for homogeneity of dispersion between pre-translocation and post-translocation invertebrate communities using the *betadisper()* function from the “vegan” package in R.

### Statistical analysis - benthic macroinvertebrate survival rates

To evaluate and compare the survival rates of benthic macroinvertebrate taxa across different sensitivity levels, we classified the 62 identified taxa into three tolerance categories: sensitive (21 taxa), moderately sensitive (27 taxa), and tolerant (14 taxa), using the Maryland Biological Stream Monitoring (MBSS) database, which categorizes different benthic macroinvertebrate taxa based on their response to organic pollution (Stribling et al. 1998). We calculated abundance per tube by summing up the abundance for selected taxa within each category. If a taxon was present in both spring and fall, we summed the spring and fall abundances. Using abundance per tube as the response variable, we fitted a Generalized Linear Model (GLM) with a negative binomial family to account for the overdispersion in the abundance data (Stoklosa et al. 2022). In the model, we included site type (pre-translocation or post-translocation) and sensitivity as fixed effects. This was executed using the *lmer()* function from the lme4 package in R (Bates et al. 2015, 2015). To evaluate how changes in benthic macroinvertebrate abundance following translocation vary across the selected watersheds, we fitted three two-factor GLMs with a negative binomial distribution, one for each watershed. In each case, we used the total abundance of one of the sensitivity categories as the response variable and site and watershed as fixed predictors.

Finally, we compared the change in abundance between pre-translocation and post-translocation tubes for individual taxa by fitting a single factor fixed-effect GLM with a negative binomial family using site type as a fixed predictor and the abundance of each taxon as the response variable. For this analysis, we only included taxa that were found in at least five donor tubes, i.e., sensitive (16), moderately sensitive (6), and tolerant (6). We repeated this for every taxon that occurred in at least five habitat tubes. Because community composition and environmental conditions differ in the spring and fall, we separated the dataset into spring and fall subsets and ran the analysis separately. We extracted model coefficients and associated statistics, i.e., effect sizes, p-values, and standard errors, using the *broom()* package in R (Robinson 2014). The extracted statistics were collated into a data frame, and 95% Confidence Intervals (CI) were calculated around the effect sizes. We used the *ggplot*() function from the ggplot2 package to create coefficient plots showing the estimated effect sizes and CIs for each taxon.

### Statistical analysis - substrate comparison

To assess the effectiveness of rock and leaf substrates in aggregating benthic macroinvertebrate communities for translocation, we compared community composition, diversity, and abundance of benthic macroinvertebrates between the two substrate types. For community composition, we separated the full dataset into fall and spring samples and analyzed each independently to avoid the pronounced seasonal difference in community composition overwhelming the treatment-related effect. We then used NMDS to ordinate rock and leaf communities in a three-dimensional solution (k=3) using Bray-Curtis distance and a maximum of 1000 iterations. We used PERMANOVA to test for a significant difference in the centroids. Before running the ordination and the PERMANOVA, we square-root transformed the raw abundance data to reduce the dominance of highly abundant taxa, and we removed rare taxa as above to avoid inflating the dissimilarity between invertebrate communities on both substrates.

After running the ordination and PERMANOVA, we tested for homogeneity of dispersion between the two groups to ensure that observed results are not driven by differences in variability within the groups.

To compare diversity across rock and leaf substrates, we used four metrics, i.e., taxa richness, EPT Richness, Shannon diversity, and abundance. We calculated taxa richness as the number of taxa in each tube, and EPT richness as the number of EPT taxa in each tube. We calculated Shannon diversity for each of the habitat tubes using the *diversity()* function from the vegan package in R, while we calculated abundance by summing up the total number of individuals of all taxa in a tube.

Preliminary two-factor Linear Models (LM) with substrate and season as fixed effects showed that the substrate X season interaction was insignificant for abundance (F = 0.3293, p = 0.56835), taxa richness (F = 0.0608, p= 0.8062), and Shannon diversity (F = 0.0374, p = 0.847), so here we dropped the season effect and fit a single factor fixed effect Linear Model (LM), using each metric as the response variable and substrate as the predictor variable. The residuals of our taxa richness, Shannon diversity, and abundance data met the normality and homogeneity of variance assumptions, so we used a Gaussian distribution. The residuals of EPT richness, however, did not meet the normality and homogeneity of variance assumptions, so we used a Poisson distribution. A preliminary two-factor GLM using substrate type and season as fixed effects showed that the substrate X season interaction was also insignificant for EPT Richness (χ^2^ = 0.1411, df = 1, p = 0.9054), so here we fit a single-factor fixed-effect GLM using EPT richness as the response variable and substrate as the predictor variable. We summarize all the linear models described above in Table 1. All data and code are available at https://github.com/Ibrahim-Fagbohun/Survival_Experiment/

**Table 1:** Summary of Statistical Models.

| Model | Model Type | Distribution | Response | Fixed Effect | Interaction |
| --- | --- | --- | --- | --- | --- |
| Model 1 | GLM | Negative binomial | Abundance | Site, Sensitivity | Site × Sensitivity |
| Model 2 | GLM | Negative binomial | Sensitive taxa abundance | Site, Watershed | Site × Watershed |
| Model 3 | GLM | Negative binomial | Tolerant taxa abundance | Site, Watershed | Site × Watershed |
| Model 4 | GLM | Negative binomial | Moderately Sensitive taxa abundance | Site, Watershed | Site × Watershed |
| Model 5 | GLM | Negative binomial | Individual taxa abundance (Fall) | Site | No interaction |
| Model 6 | GLM | Negative binomial | Individual taxa abundance (Spring) | Site | No interaction |
| Model 7 | LM | Gaussian | Taxa Richness | Substrate Type | No interaction |
| Model 8 | LM | Gaussian | Shannon Diversity | Substrate Type | No interaction |
| Model 9 | LM | Gaussian | Abundance | Substrate Type | No interaction |
| Model 10 | GLM | Poisson | EPT Richness | Substrate Type | No interaction |

## Results

### Community composition (pre-translocation vs post-translocation)

Using a three-dimensional solution (k =3), the NMDS ordination for the fall 2024 and spring 2025 community compositions both reached convergence after 20 iterations with stress values of 0.15 and 0.16, respectively. The PERMANOVA analysis showed a significant change in community composition between pre-translocation and post-translocation tubes both in fall 2024 (*F* = 7.6626, *p* = 0.001, *R^2^* = 0.1185, df = 57, Table S1) and spring 2025 (*F* = 14.977, *p* = 0.001, *R^2^* = 0.2052, df = 58, Table S2) suggesting a change in community composition following invertebrate translocation to the restored streams in both seasons. The homogeneity of dispersion test confirmed similar dispersion between both groups in the fall (*F* = 3.5959, *p* = 0.063, df = 57, Table S3; Figure 2) and spring (*F* = 1.4626, *p* = 0.2314, df = 58, Table S4; Figure 3), indicating that the observed differences can be confidently attributed to differences in centroids.

**Figure 2:**
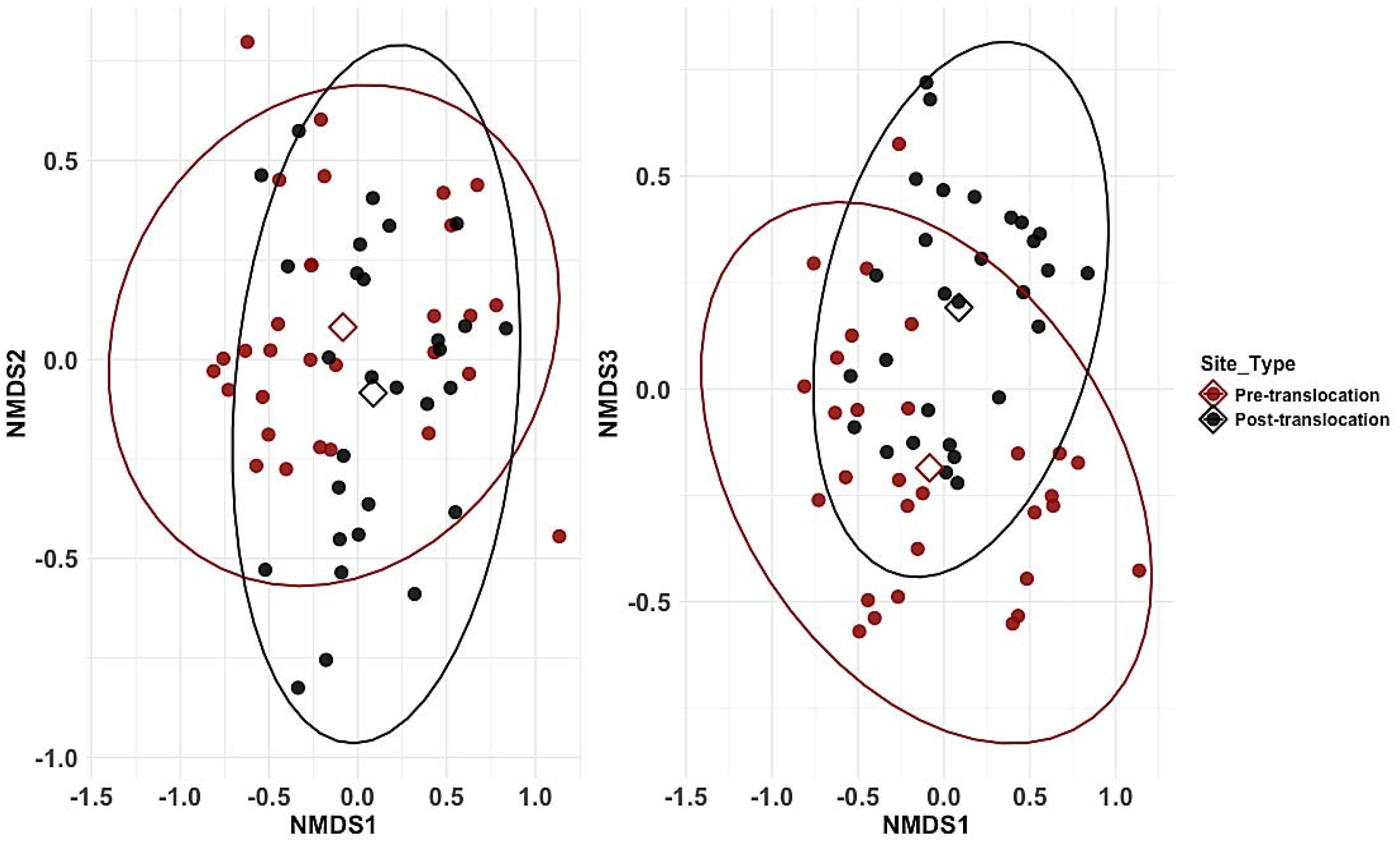
Nonmetric multidimensional scaling (NMDS) ordination of macroinvertebrate communities based on Bray-Curtis dissimilarities in pre-translocation and post-translocation tubes for Fall 2024 samples. The left plot shows axes 1 vs axes 2, while the right plot shows axes 1 vs axes 3. The open diamond shapes represent the centroids for the pre-translocation and post-translocation groups.

**Figure 3:**
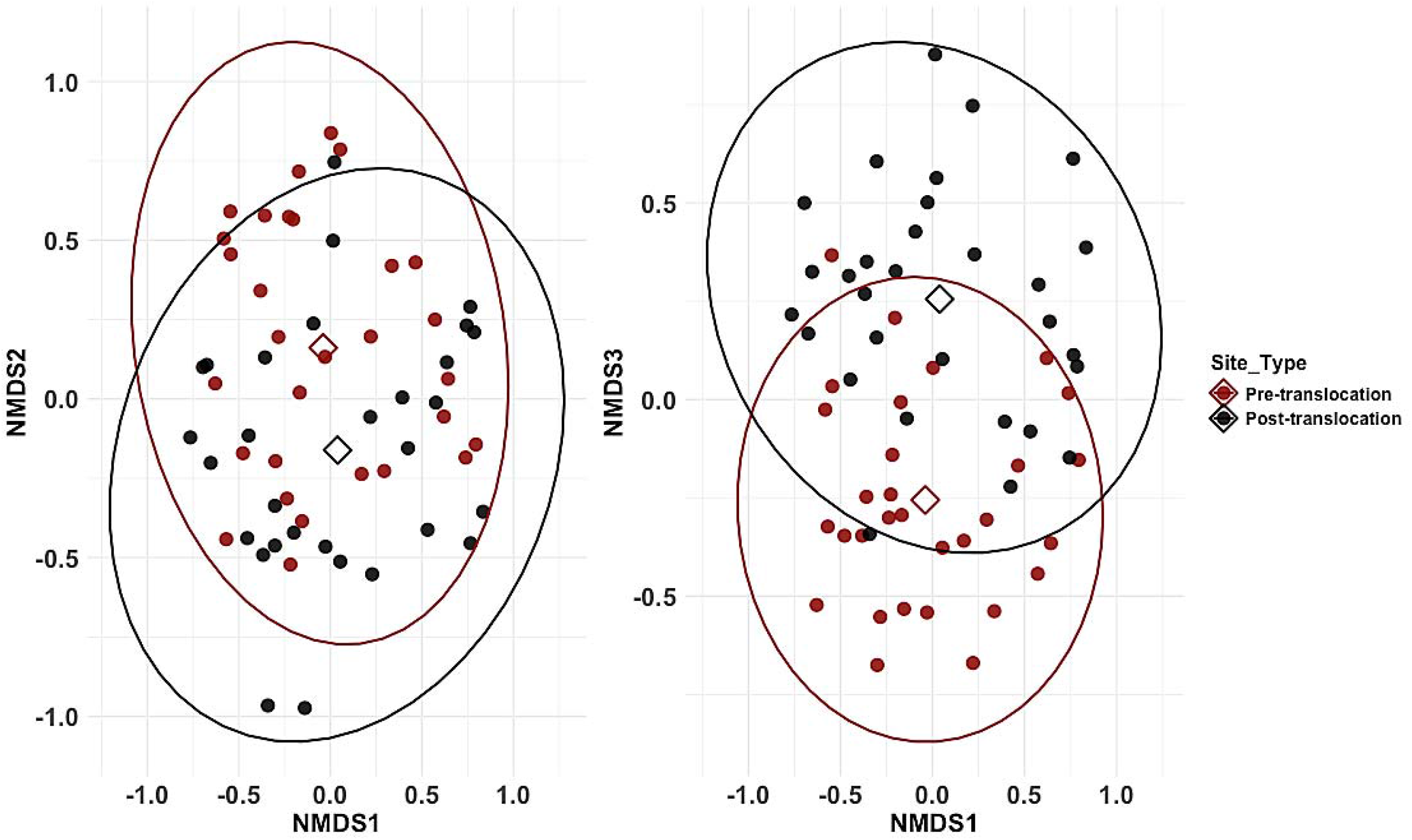
Nonmetric multidimensional scaling (NMDS) ordination of macroinvertebrate communities based on Bray-Curtis dissimilarities in pre-translocation and post-translocation tubes for Spring 2025 samples. The left plot shows axes 1 vs axes 2, while the right plot shows axes 1 vs axes 3. The open diamond shapes represent the centroids for the pre-translocation and post-translocatio groups.

### Benthic macroinvertebrate survival

The two-factor GLM (Model 1, Table S5) revealed a general decline in the abundance of benthic macroinvertebrates across all sensitivity categories (Figure 4). Although the interaction test revealed a marginal interaction between site and sensitivity (Wald = 4.72, p = 0.095, df = 2, Table S6), the follow-up simple effects pairwise contrast using emmeans showed that the magnitude of decline following translocation varied among sensitivity categories (Table S7).

**Figure 4:**
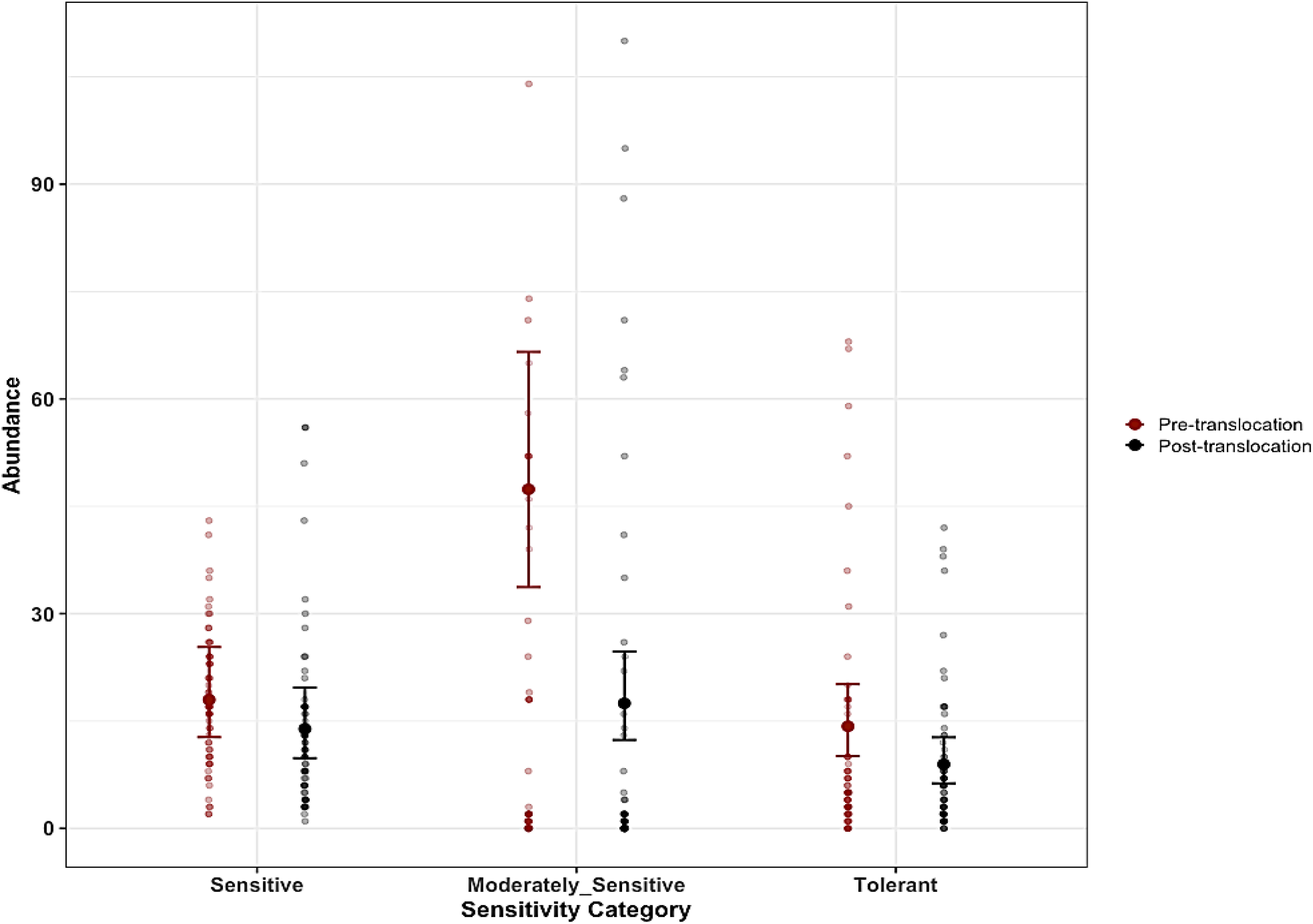
Mean plots showing benthic macroinvertebrate abundance across sensitivity categories pre- and post-translocation. Points show means ± 95% CI, with jittered raw data. Negative binomial GLM revealed significant main effects of site (*x*² = 16.19, p= <0.001) and sensitivity (*x*² = 26.6, p= <0.001), and no site * sensitivity interaction (*x*² = 4.72, p = 0.095). Letters indicate pairwise statistical differences within each sensitivity category: groups with the same letter are not statistically different.

Moderately sensitive taxa showed a strong and significant abundance reduction following translocation (*b* = −0.997, p < 0.0001), corresponding to approximately 63% decline in abundance. Tolerant taxa showed a 37% decline (*b* = −0.466, p = 0.063) while sensitive taxa showed a 23% decline (*b* = −0.257, p =0.3035).

The decline observed following translocations was inconsistent across watersheds. For sensitive taxa, the two-factor GLM (Model 2, Table S8), including site and watershed as fixed effects, showed a significant interaction between site and watershed (Wald = 11.831, p = 0.003, df = 2, Table S9). Follow-up simple effect pairwise contrasts revealed a relatively large and significant decline in sensitive taxa abundance at both Little Patuxent (*b* = −0.626, p < 0.001) and Potomac River (*b* = −0.571, p = 0.0034) watersheds (Table S10), while abundance change at the Loch Raven watershed was minimal and not statistically significant (*b* = 0.168, p = 0.3464; Figure 5a).

**Figure 5:**
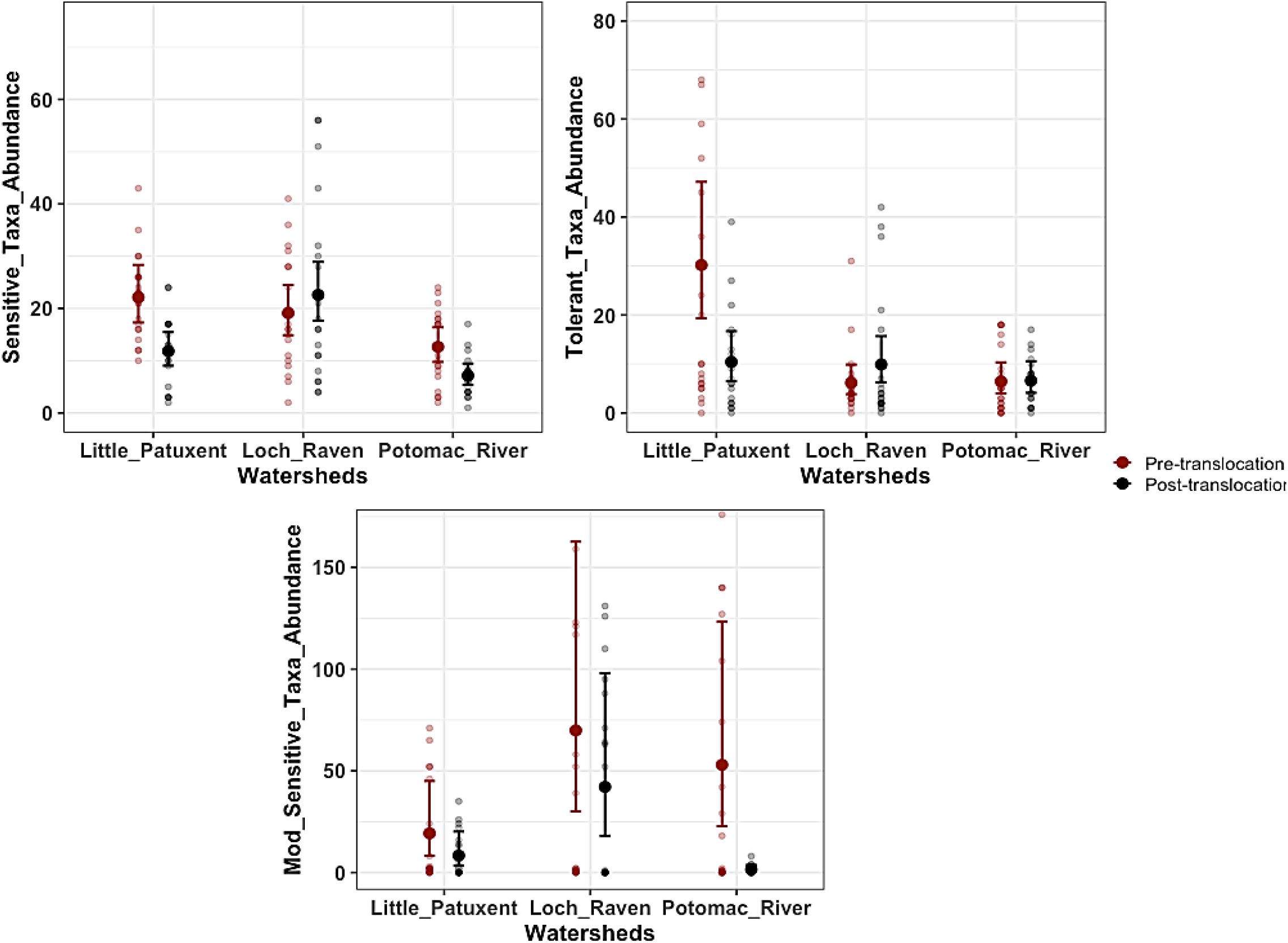
Mean benthic macroinvertebrate abundance (±95% CI) across watersheds pre- and post-translocation for sensitive, moderately sensitive, and tolerant taxa. Jittered points represent raw abundance values. For sensitive taxa, a negative binomial GLM detected significant effects of site (² = 11.36, p < 0.001) and a site × watershed interaction (² = 11.83, p = 0.002). For tolerant taxa, both the site effect (*x*² = 10.34, p = 0.001) and the site × watershed interaction (*x*² = 11.28, p = 0.003) were significant. For moderately sensitive taxa, the site effect was not significant (*x*² = 1.79, p = 0.18), but the site × watershed interaction was significant (*x*² = 14.27, p < 0.001). Letters indicate pairwise statistical differences within each sensitivity category: groups with the same letter are not statistically different.

For tolerant taxa, the two-factor GLM (Model 3, Table S11) also showed a significant site and watershed interaction on tolerant macroinvertebrate abundance (Wald = 11.275, p = 0.003, df = 2, Table S12). The follow-up simple effect contrast reveals a large and significant abundance decline in the Little Patuxent watershed (*b* = −1.064, p = 0.0013). Unlike Little Patuxent, Potomac River and Loch Raven watersheds showed little and non-significant abundance change post-translocation (Figure 5b, Table S13).

For moderately sensitive taxa, the two-factor GLM (Model 4, Table S14) also showed a significant interaction of site and watershed on invertebrate abundance (Wald = 14.27, p < 0.001, df = 2, Table S15). Although there was a general decline in abundance for post-translocation tubes across all watersheds, the decline in the Potomac River watershed was relatively large and significant (*b* = −3.564, p < 0.0001) compared to Little Patuxent (*b* = −0.833, p = 0.1814) and Loch Raven (*b* = −0.507, p = 0.4056) Watersheds (Figure 5c, Table S16).

Taxon-level models (Models 5 and 6) comparing the abundance of each taxon pre-translocation and post-translocation revealed substantial variation among taxa irrespective of their sensitivity category. For instance, 67% of the sensitive taxa in the fall, including *Isonychia, Ameletus, Diplectrona, Stenonema (Maccaffertium), Nigronia*, and *Rhyacophila,* did not have estimated abundance effect sizes that differed significantly from zero (Figure 6, Table S17).

**Figure 6:**
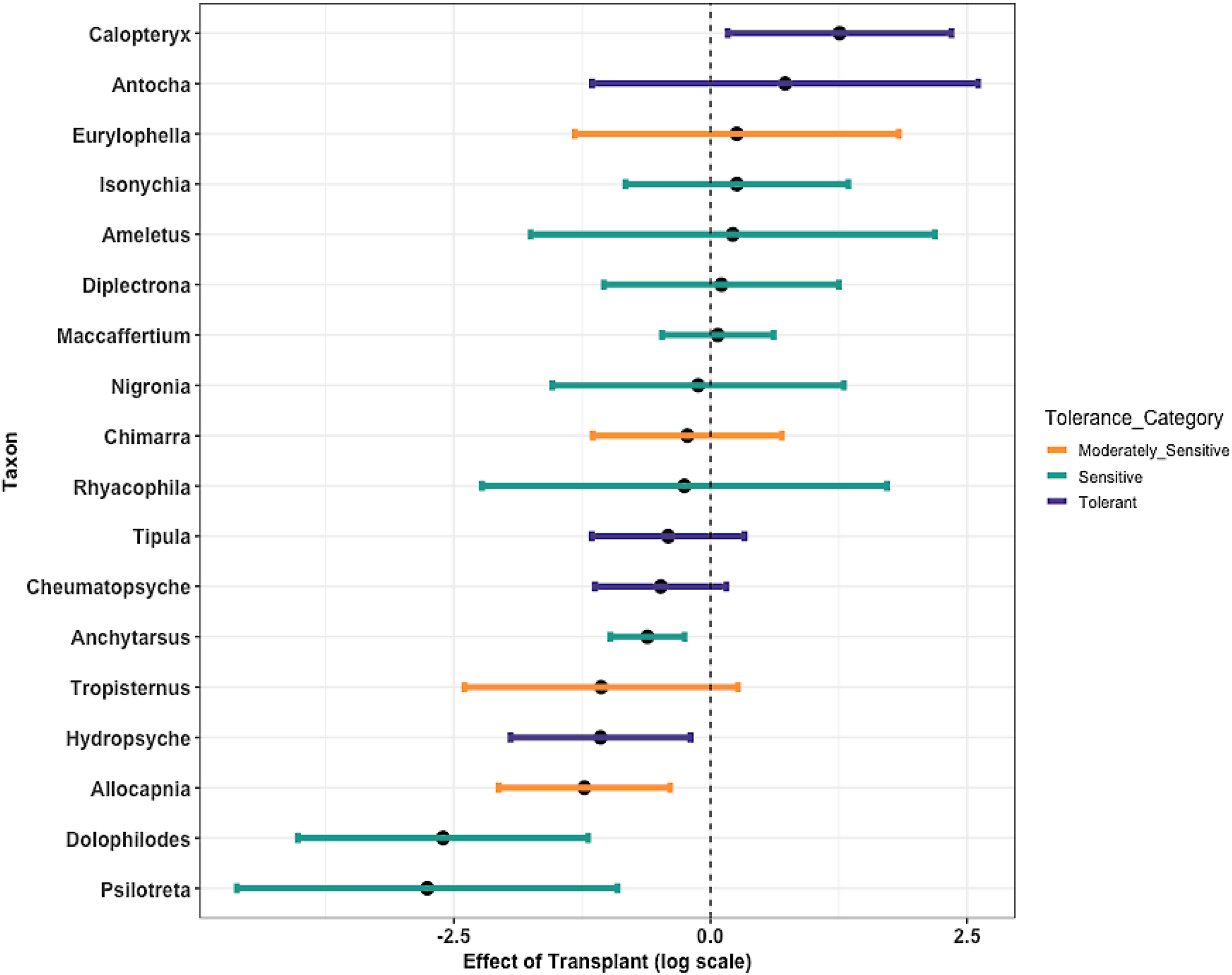
Coefficient plots summarizing taxon-specific GLM results for all study sites from Fall 2024 sampling. Black dots show effect sizes representing the log-scale difference in taxa abundance between pre- and post-translocation, with negative values indicating abundance decline post-translocation and vice versa. Whiskers represent 95% CI. All CI’s overlapping zero are not significant at p < 0.05.

However, other sensitive taxa, including *Anchytarsus, Dolophilodes, and Psilotreta,* all showed significant negative abundance effect sizes post-translocation (Figure 6, Table S17). In the spring, 61% of sensitive taxa, including *Stenonema* (*Maccaffertium), Polycentropus, Nigronia, Timpanoga, Plectrocnemia, Penelomax, Isonychia,* and *Anchytarsus,* did not have abundance effect sizes significantly different from zero, while 39% were significantly lower than zero (Figure 7, Table S18). For moderately sensitive taxa, 75% and 66% of taxa had abundance effect sizes not significantly different from zero in the fall and spring, respectively. For the tolerant category, 80% and 100% of taxa in the fall and spring, respectively, had no significant change in abundance.

**Figure 7:**
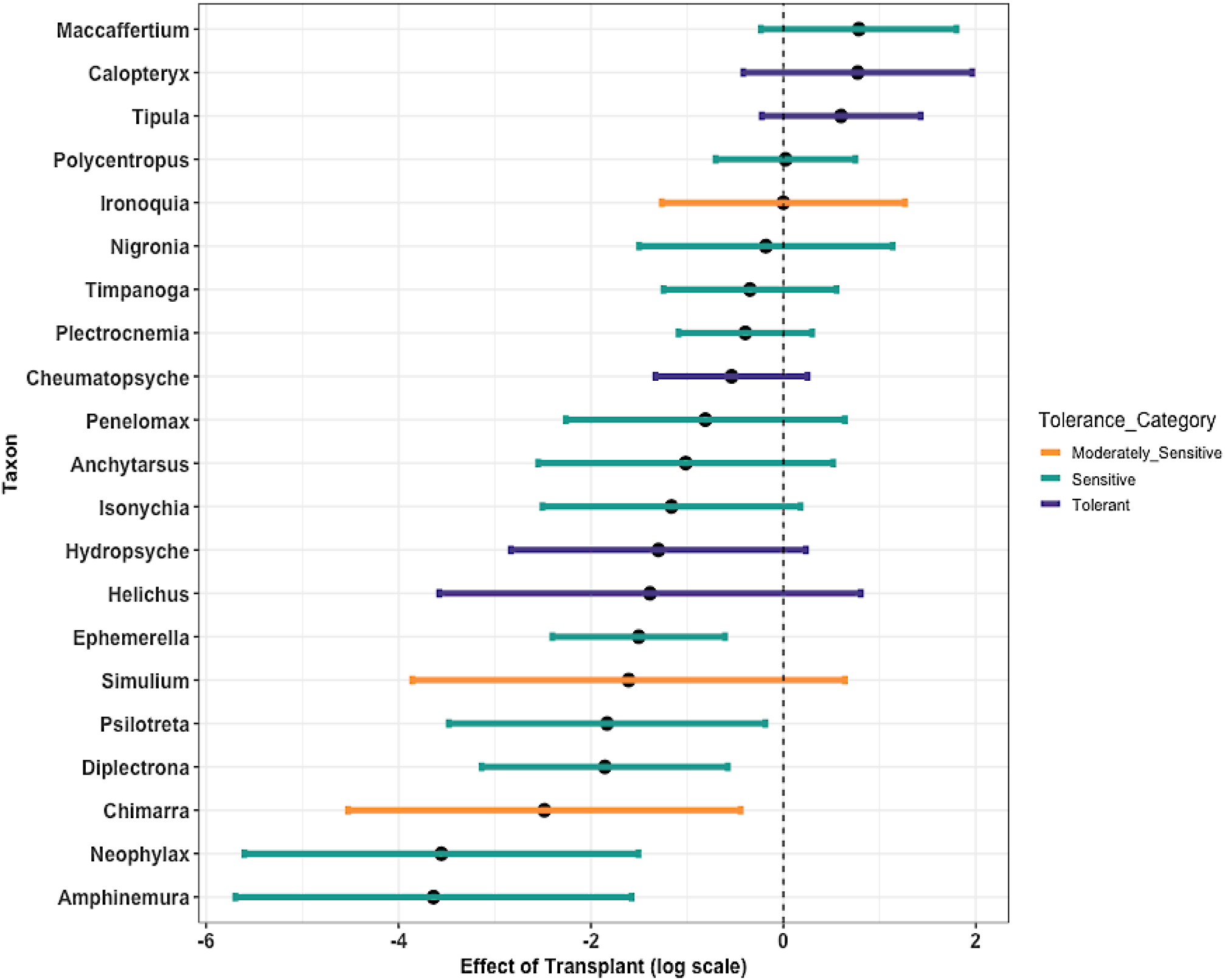
Coefficient plots summarizing taxon-specific GLM results for all study sites from spring 20025 sampling. Black dots show effect sizes representing the log-scale difference in taxa abundance between pre- and post-translocation, with negative values indicating abundance decline post-translocation and vice versa. Whiskers represent 95% CI. All CI’s overlapping zero are not significant at p < 0.05.

### Substrate analysis

The NMDS ordinations comparing community composition between leaf and rock substrates in fall 2024 and spring 2025 converged after 20 iterations, with stress values of 0.11 and 0.12, respectively (Figures 8 and 9). The PERMANOVA analysis showed no significant difference in centroids based on the calculated Bray-Curtis distance between the leaf and the rock benthic macroinvertebrate communities in the fall (F = 1.9371, p = 0.107, R^2^ = 0.0647, df = 28, Table S19) and spring (F = 1.5663, p = 0.144, R^2^ = 0.05, df = 28, Table S20). Similarly, the linear models (Models 7,8, and 9) comparing taxa richness, EPT richness, and Shannon diversity, between substrates revealed that rock and leaf substrates have similar Taxa richness (*b* = −0.8, p = 0.3868), EPT richness (*b* = 0.05, p = 0.713) and Shannon diversity (*b* = 0.16, p = 0.27), and the similarity stays consistent across the fall and spring seasons as indicated by a non-significant substrate - season interaction term for all three diversity metrics. Despite no effect of substrate type on diversity, the linear model (Model 10) with abundance as a response variable showed that leaf substrates contained a significantly higher number of macroinvertebrate individuals than rock substrates (*b* = −45.00, p = 0.027; Table S23, Figure 10).

**Figure 8:**
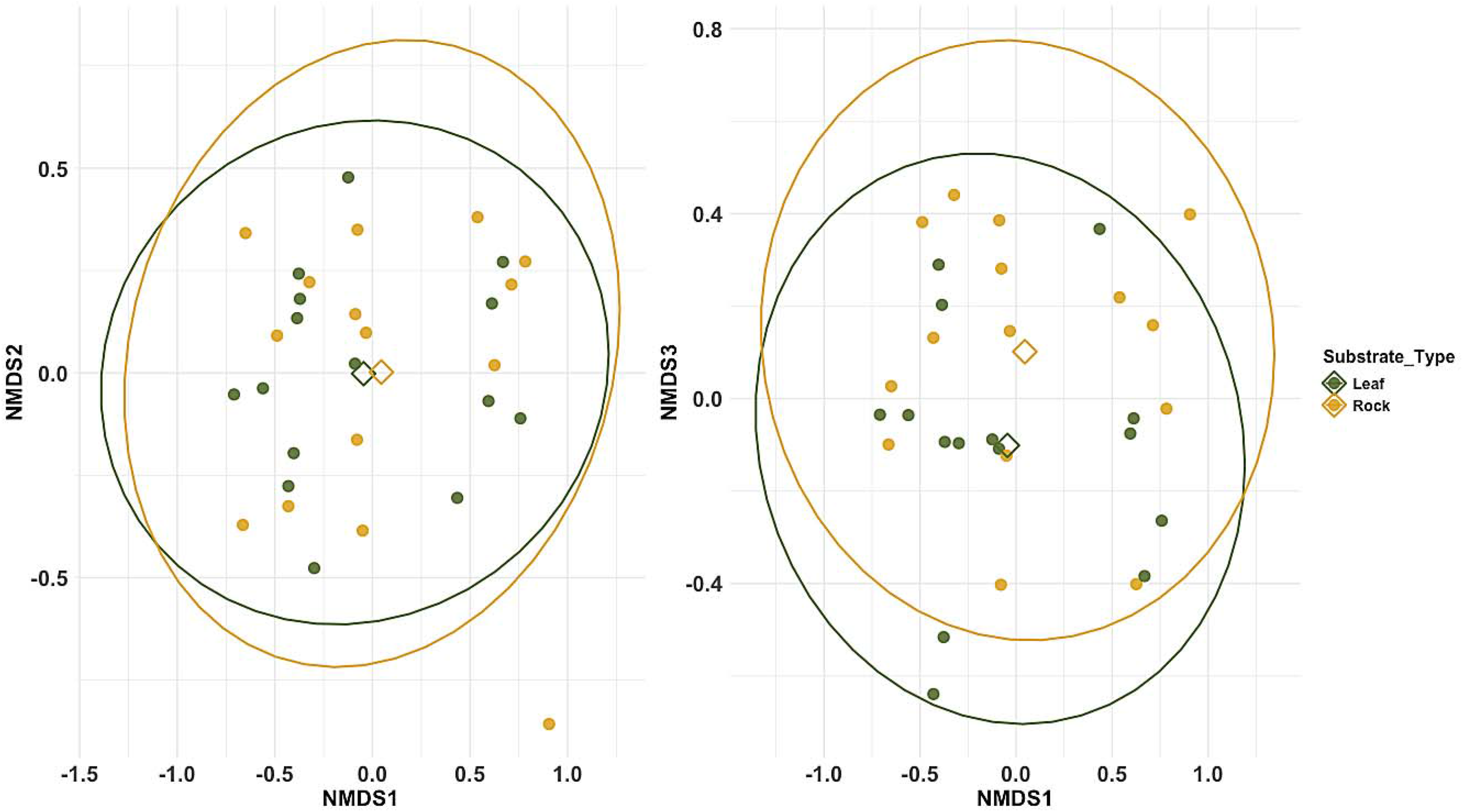
Bray-Curtis-based NMDS showing no significant differences between benthic macroinvertebrate communities on the leaf and rock substrates across donor sites in fall 2024. Individual points (circles) represent each habitat tube, while the open diamond represents the centroids for each substrate.

**Figure 9:**
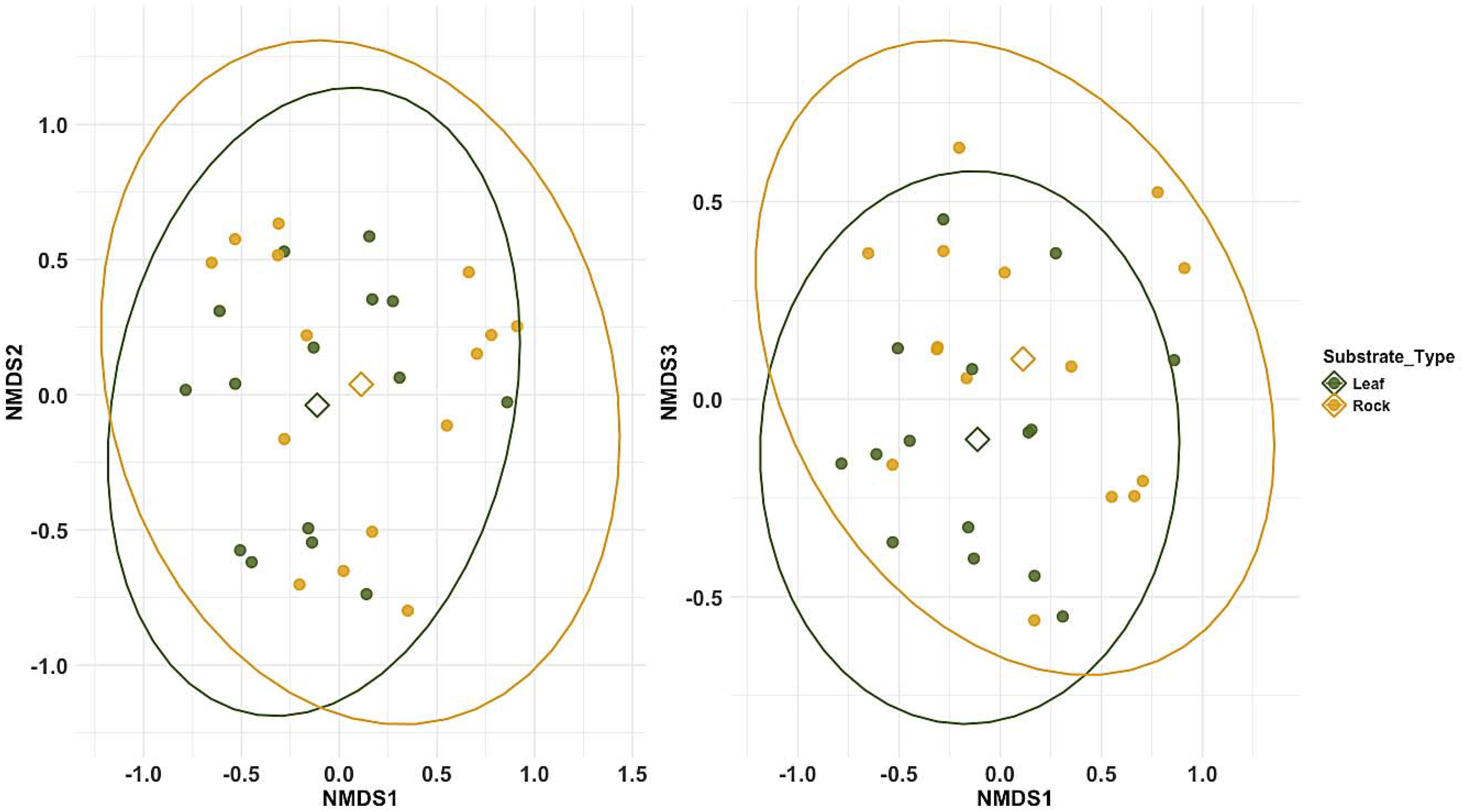
Bray-Curtis based NMDS showing no significant differences between benthic macroinvertebrate communities on the leaf and rock substrates across donor sites in fall 2024. Individual points (circles) represent each habitat tube, while the open diamond represents the centroids for each substrate.

**Figure 10:**
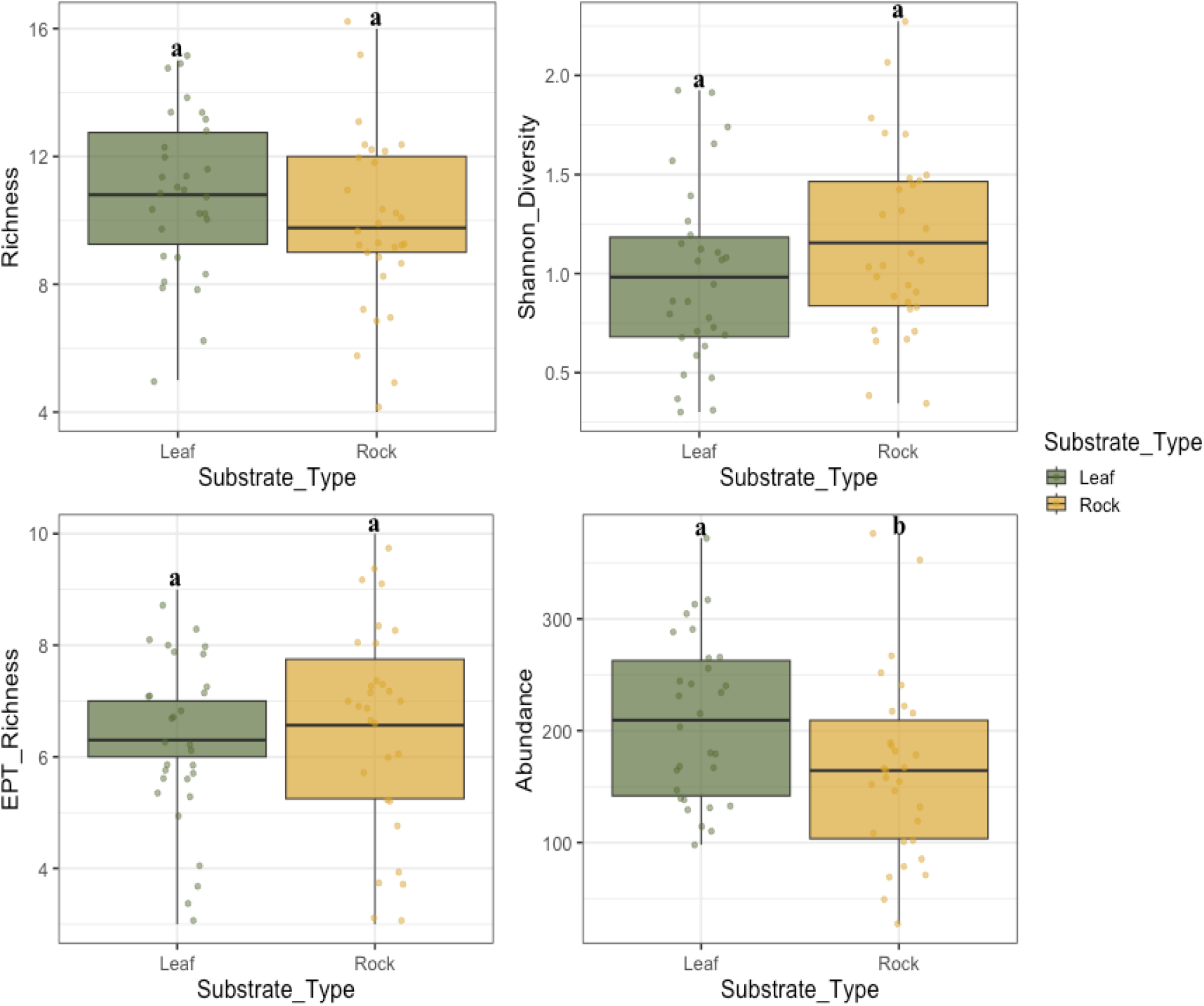
Boxplots showing the effect of substrate on different metrics of benthic macroinvertebrate diversity and abundance after four weeks of colonization in the donor streams in fall 2024 and spring 2025. Boxes summarize diversity estimates as the mean, 25th and 75thperceantiles, and minimum and maximum values. Letters indicate pairwise statistical differences within each metric: groups with the same letter are not statistically different.

## Discussion

The lack of improvement in benthic macroinvertebrate composition, diversity, and abundance after stream restorations has been widely reported (Palmer et al. 2010; Louhi et al. 2011; Stranko et al. 2012; Leps et al. 2016; Hilderbrand & Acord 2019). These studies have strengthened the position that current and common restoration practices often do not improve stream conditions sufficiently to support higher macroinvertebrate diversity (Haase et al. 2013a; Hörchner et al. 2024). Our results suggest that translocating benthic macroinvertebrates from donor streams has the potential to assist recovery of macroinvertebrate communities in stream restorations. Although our experiment lasted four weeks, taxa representing all sensitivity levels survived during both fall and spring translocations. This suggests that restored streams may be able to support more macroinvertebrate diversity than they currently do.

After harvesting macroinvertebrates from reference streams and leaving them in restored streams for 28 days, we observed a general decline in the abundance of macroinvertebrates across all sensitivity categories. We expected some decline given that biotic interactions, such as increased competition and predation, would occur in a closed system. Abiotic processes could have led to declines as well: reduced substrate and food quality, reduced flow velocity, and sediment accumulation within the habitat tubes could cause general mortality across all sensitivity levels. However, we predicted that if water quality conditions within the restored stream are inhospitable to sensitive taxa, we would observe a disproportionately larger decline in the abundance of sensitive macroinvertebrate taxa relative to moderately sensitive and tolerant taxa. Our results showed otherwise: sensitive macroinvertebrate taxa declined by only 23%, lower than the 37% observed for tolerant taxa, though still significantly lower than the 63% decline reported for moderately sensitive taxa. Overall, these results support our initial hypothesis that sensitive benthic macroinvertebrates can survive in restored streams.

Comparing the rate of sensitive taxa decline across watersheds, we observed lower rates of abundance decline in the restored sites in the Loch Raven reservoir and Potomac River watersheds, with a relatively large decline in the restored site of the Little Patuxent River watershed. A similar pattern was observed for tolerant taxa, suggesting variation in the habitat quality of restored streams, and thus the survivability of translocated taxa, might differ among restored streams. This mechanism underlying this pattern is not clear to us, but may depend on a suite of factors, including land use, age of restoration, type of restoration, impervious area, thermal regime, flow regime, and food resources; all of which contribute to overall habitat quality (Allan 2004; Hughes 2007; Thompson et al. 2018b).

Focusing on individual taxa performance following translocations, we found that 11 out of 16 sensitive macroinvertebrate taxa, including *Isonychia, Ameletus, Diplectrona, Stenonema (Maccaffertium), Nigronia*, *Rhyacophila, Polycentropus, Timpanoga, Plectrocnemia, Penelomax,* and *Anchytarsus,* survived in restored streams. We also found that 5 out of 6 moderately sensitive macroinvertebrate taxa, including *Eurylophella*, *Chimarra, Tropisternus, Ironoquia*, and *Simulium*, survived in restored streams. Similar findings of sensitive and moderately sensitive benthic macroinvertebrate taxa surviving and even persisting in restored streams following translocations into streams where they have been previously extirpated have also been reported in Washington, USA (Macneale 2020) and the Elsebach River in Germany (Gellert et al. 2025).

Our results suggest that stream restorations may be able to support sensitive benthic macroinvertebrate taxa currently absent, and the generally poor biological condition of restored streams cannot be attributed solely to habitat or water quality. Sensitive taxa survived when experimentally translocated into restored streams, demonstrating that they can survive in these streams, at least in the short term. Our findings suggest dispersal limitation as a possible key constraint on the natural recovery of sensitive macroinvertebrate communities (Patrick et al. 2021). The inability of benthic macroinvertebrates to disperse to and colonize channels beyond 5 km of their source population has been previously indicated (Lake et al. 2007; Sundermann et al. 2011; Stoll et al. 2016). Dispersal can constrain the recovery of macroinvertebrates in restored streams if sensitive species are either unable to reach restored reaches or if they do so in very small numbers, making them highly vulnerable to demographic, environmental, and genetic stochasticity, which can prevent the establishment of a self-sustaining population (Parkyn & Smith 2011). To bridge this dispersal gap, whole-community translocation has been recommended as a relatively simple, effective, and inexpensive method, though it remains experimentally untested (Haase & Pilotto 2019; Contos et al. 2021). Although Jourdan et al. (2019) argued that such translocation might increase the risk of pathogen and invasive species transport to restored sites, here we minimized these risks by pairing recipient sites with donor streams within the same watershed or adjacent watersheds and following a standard gear decontamination protocol when moving between sites.

We also tested the effectiveness of two substrates, oak leaves and river rocks, in translocating macroinvertebrate communities. Our results showed that both substrates performed similarly in accumulating a diverse community of benthic macroinvertebrates. Both substrates showed similar taxa richness, EPT richness, and Shannon diversity, indicating that either substrate can be valuable for whole invertebrate community translocation. However, leaf tubes on average accumulated 27% more individuals than rock tubes. Leaf substrates accumulating a relatively higher abundance of invertebrates have also been reported in Dumeier et al. (2018), where they showed that a mixture of wood and beech leaf substrate accumulated a relatively more diverse and abundant macroinvertebrate community than other substrates, including gravel, wood, and alder leaf. Considering the importance of abundance in ensuring the survival and continuity of reintroduced species, we recommend the prioritization of leaf substrates for whole community translocations.

In conclusion, our results demonstrate that the limited recovery of sensitive macroinvertebrate taxa in restored streams may result as much from dispersal constraints as from habitat limitations. By experimentally removing dispersal barriers, our results suggest that a whole community translocation using passive exposure of rock and leaf substrates in reference streams for 4-6 weeks can aggregate a morphologically and functionally diverse and abundant community of macroinvertebrates that can be translocated to restored streams. Collectively, these findings identify whole-community translocations as a promising tool for overcoming dispersal barriers, accelerating biodiversity recovery, and improving biological condition in restored streams

## Acknowledgements

Funding for this project was provided by the Chesapeake Bay Trust Award 22062. Additional support to IRF was provided by a Huck Institute of Life Sciences Graduate Fellowship from Penn State University. Additional support to DCA was provided by the USDA National Institute of Food and Agriculture and Hatch Appropriations under Project #PEN04817 and Accession #7003724, and to JNS under Project PEN04810 and Accession #7003691.

