## Supplemental Figures and Tables. for "Whole-community translocation shows high survival potential of pollution-sensitive benthic macroinvertebrate taxa in restored streams"

**Supplementary Material**

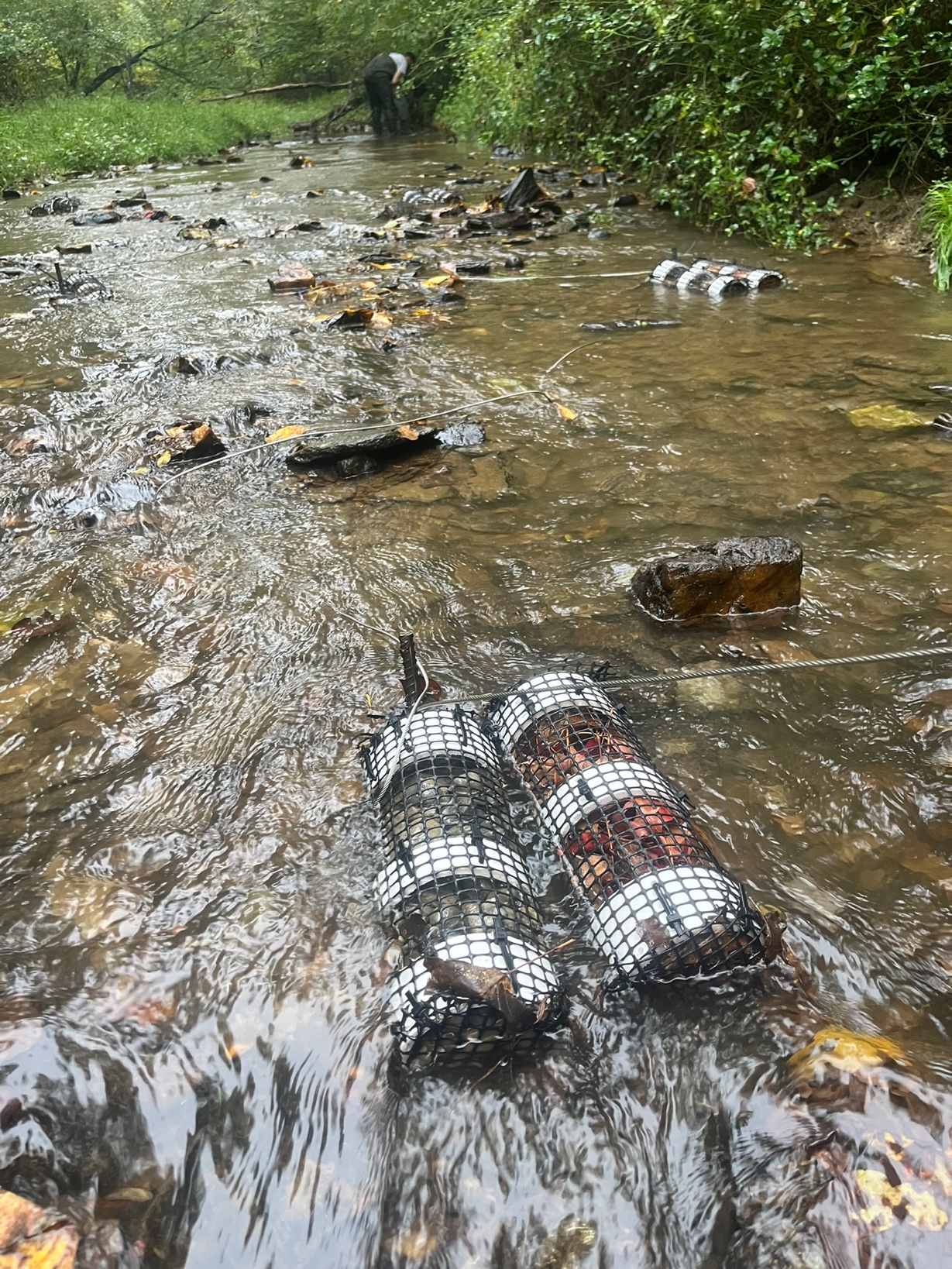

Figure S1: Arrangement of paired habitat tubes in the donor sites. Each stake held one rock tube, and one leaf tube positioned side-by-side to standardize exposure. Each donor site contained 10 stakes, supporting a total of 20 habitat tubes (10 rocks and 10 leaves) deployed for four weeks to ensure enough time for macroinvertebrate colonization.

Table S1: PERMANOVA tests evaluating the effects of restored streams on macroinvertebrate community composition in Fall 2024 across all donor sites.

| Source | df | Sum of Squares | R^2^ | F | p-value |
| --- | --- | --- | --- | --- | --- |
| Site type | 1 | 0.7266 | 0.1185 | 7.6626 | 0.001 |
| Residuals | 57 | 5.4048 | 0.8815 |  |  |
| Total | 58 | 6.1314 | 1.00 |  |  |

Table S2: PERMANOVA tests evaluating the effects of restored streams on macroinvertebrate community composition in Spring 2025 across all donor sites.

| Source | df | Sum of Squares | R^2^ | F | p-value |
| --- | --- | --- | --- | --- | --- |
| Site type | 1 | 1.6369 | 0.2052 | 14.977 | 0.001 |
| Residuals | 58 | 6.3391 | 0.7947 |  |  |
| Total | 59 | 7.9760 | 1.00 |  |  |

Table S3: PERMDISP results assessing homogeneity of multivariate dispersion between pre- and post-translocation macroinvertebrate communities across donor sites in Fall 2024.

| Source | df | Sum of Squares | Mean Square | F | p-value |
| --- | --- | --- | --- | --- | --- |
| Site Type | 1 | 0.02683 | 0.0268 | 3.5959 | 0.063 |
| Residuals | 57 | 0.425 | 0.00746 |  |  |

Table S4: PERMDISP results assessing homogeneity of multivariate dispersion between pre- and post-translocation macroinvertebrate communities across donor sites in Fall 2024.

| Source | df | Sum of Squares | Mean Square | F | p-value |
| --- | --- | --- | --- | --- | --- |
| Site Type | 1 | 0.00850 | 0.00850 | 1.4626 | 0.2314 |
| Residuals | 58 | 0.33714 | 0.00746 | 0.0058 |  |

Table S5: Coefficient estimates from a negative binomial GLM estimating the effect of site, sensitivity, and their interaction on macroinvertebrate abundance. The table reports parameter estimates relative to reference levels, SE, test statistic, and p-values.

| Term | Estimate | SE | z | p-value |
| --- | --- | --- | --- | --- |
| Site type (Post-Translocation) | 0.9971 | 0.25 | 4.02 | <0.001 |
| Sensitivity (Sensitive) | -0.2290 | 0.25 | -0.914 | <0.001 |
| Sensitivity (Tolerant) | -0.6692 | 0.25 | -2.656 | <0.001 |
| Post-Translocation × Sensitive | -0.74 | 0.35 | -2.105 | 0.03 |
| Post-Translocation × Tolerant | -0.53 | 0.35 | -1.504 | 0.13 |

Table S6: Wald $\chi^{2}$ tests from negative binomial GLM evaluating the effects of site, sensitivity, and their interaction on macroinvertebrate abundance.

| Effect | $\chi^{2}$ | df | p-value |
| --- | --- | --- | --- |
| Site | 15.97 | 1 | < 0.001 |
| Sensitivity | 26.21 | 2 | < 0.001 |
| Site × Sensitivity | 4.72 | 2 | 0.095 |

Table S7: Emmeans simple effect contrast (post-translocation vs pre-translocation) within each sensitivity category for the negative binomial GLM. All estimates are on the log scale.

| Contrast | Estimates | SE | z-ratio | p-value |
| --- | --- | --- | --- | --- |
| Sensitive | -0.257 | 0.250 | -1.029 | 0.3035 |
| Moderately sensitive | -0.997 | 0.248 | -4.023 | <0.0001 |
| Tolerant | -0.466 | 0.251 | -1.856 | 0.0635 |

Table S8: Coefficient estimates from negative binomial GLM testing the effect of site, watershed, and their interaction on sensitive macroinvertebrate abundance.

| Term | Estimate | SE | z | p-value |
| --- | --- | --- | --- | --- |
| Site type (Post-translocation) | 0.63 | 0.18 | 3.37 | < 0.001 |
| Watershed (Loch Raven) | 0.64 | 0.18 | 3.481 | <0.001 |
| Watershed (Potomac River) | -0.50 | 0.19 | -2.547 | 0.01 |
| Pre-Translocation × Loch Raven | -0.79 | 0.26 | -3.082 | 0.002 |
| Pre -Translocation × Potomac River | -0.06 | 0.27 | -0.207 | 0.84 |

Table S9: Wald $\chi^{2}$ tests from negative binomial GLM evaluating the effects of site, watershed, and their interaction on sensitive macroinvertebrate taxa abundance

| Effect | $\chi^{2}$ | df | p-value |
| --- | --- | --- | --- |
| Site | 11.359 | 1 | < 0.001 |
| Watershed | 37.256 | 2 | < 0.001 |
| Site × Watershed | 11.831 | 2 | 0.002 |

Table S10: Emmeans simple effect contrast (post-translocation vs pre-translocation) within each watershed for the negative binomial GLM, estimating the effects of site and watershed on sensitive macroinvertebrate abundance. All estimates are on the log scale.

| Contrast | Estimates | SE | z-ratio | p-value |
| --- | --- | --- | --- | --- |
| Little Patuxent | -0.626 | 0.186 | -3.370 | <0.001 |
| Loch Raven | 0.168 | 0.179 | 0.942 | 0.3464 |
| Potomac River | -0.571 | 0.195 | -2.925 | 0.0034 |

Table S11: Coefficient estimates from negative binomial GLM testing the effect of site, watershed, and their interaction on tolerant macroinvertebrate abundance

| Term | Estimate | SE | z | p-value |
| --- | --- | --- | --- | --- |
| Site type (Post-translocation) | -1.06 | 0.33 | 3.21 | 0.001 |
| Watershed (Loch Raven) | -0.05 | 0.33 | -0.153 | 0.878 |
| Watershed (Potomac River) | -0.46 | 0.33 | -1.344 | 0.1789 |
| Pre-Translocation × Loch Raven | -1.54 | 0.47 | -3.262 | 0.0011 |
| Pre-Translocation × Potomac River | -1.08 | 0.47 | -2.29 | 0.022 |

Table S12: Wald $\chi^{2}$ tests from negative binomial GLM evaluating the effects of site, watershed, and their interaction on tolerant macroinvertebrate taxa abundance

| **Effect** | $\boldsymbol{\chi}^{\boldsymbol{2}}$ | **df** | **p-value** |
| --- | --- | --- | --- |
| Site | 1.0618 | 1 | 0.001 |
| Watershed | 21.3395 | 2 | 0.336 |
| Site × Watershed | 11.2754 | 2 | 0.003 |

Table S13: Emmeans simple effect contrast (post-translocation vs pre-translocation) within each watershed for the negative binomial GLM, estimating the effects of site and watershed on tolerant macroinvertebrate abundance. All estimates are on the log scale.

| Contrast | Estimates | SE | z-ratio | p-value |
| --- | --- | --- | --- | --- |
| Little Patuxent | -1.064 | 0.331 | -3.215 | 0.0013 |
| Loch Raven | 0.476 | 0.337 | 1.414 | 0.1574 |
| Potomac River | 0.023 | 0.340 | 0.068 | 0.9461 |

Table S14: Coefficient estimates from negative binomial GLM testing the effect of site, watershed, and their interaction on moderately sensitive macroinvertebrate abundance

| Term | Estimate | SE | z | p-value |
| --- | --- | --- | --- | --- |
| Site type (Post-translocation) | 0.83 | 0.62 | 1.34 | 0.18 |
| Watershed (Loch Raven) | 1.61 | 0.62 | 2.59 | 0.009 |
| Watershed (Potomac River) | -1.71 | 0.64 | -2.655 | 0.008 |
| Pre-Translocation × Loch Raven | -0.33 | 0.87 | -0.374 | 0.71 |
| Pre-Translocation × Potomac River | 2.7 | 0.89 | 3.067 | 0.002 |

Table S15: Wald $\chi^{2}$ tests from negative binomial GLM evaluating the effects of site, watershed, and their interaction on moderately sensitive macroinvertebrate taxa abundance

| Effect | $\chi^{2}$ | df | p-value |
| --- | --- | --- | --- |
| Site | 19.637 | 1 | <0.001 |
| Watershed | 18.180 | 2 | < 0.001 |
| Site × Watershed | 14.2716 | 2 | < 0.001 |

Table S16: Emmeans simple effect contrast (post-translocation vs pre-translocation) within each watershed for the negative binomial GLM, estimating the effects of site and watershed on moderately sensitive macroinvertebrate abundance. All estimates are on the log scale.

| Contrast | Estimates | SE | z-ratio | p-value |
| --- | --- | --- | --- | --- |
| Little Patuxent | -0.833 | 0.623 | -1.337 | 0.1814 |
| Loch Raven | -0.507 | 0.610 | -0.832 | 0.4056 |
| Potomac River | -3.564 | 0.636 | -5.606 | <0.0001 |

Table S17: Results of Taxon-specific negative binomial GLMs testing the effect of restored sites on macroinvertebrate abundance in Fall 2024. Negative estimates indicate lower abundance post-translocation, while positive estimates represent higher abundance post-translocation. Models were fit separately for each taxon using a negative binomial error distribution to account for overdispersion.

| **Taxon** | **Estimate** | **SE** | **z (statistic)** | **p-value** |
| --- | --- | --- | --- | --- |
| *Allocapnia* | -1.23 | 0.426 | -2.88 | 0.003 |
| *Ameletus* | 0.22 | 1.00 | 0.21 | 0.829 |
| *Anchytarsus* | -0.61 | 0.18 | -3.33 | 0.0008 |
| *Antocha* | 0.72 | 0.96 | 0.76 | 0.448 |
| *Calopteryx* | 1.26 | 0.55 | 2.26 | 0.02 |
| *Cheumatopsyche* | -0.48 | 0.33 | -1.48 | 0.137 |
| *Chimarra* | -0.22 | 0.47 | -0.48 | 0.63 |
| *Diplectrona* | 0.11 | 0.58 | 0.18 | 0.85 |
| *Dolophilodes* | -2.60 | 0.72 | -3.61 | 0.0003 |
| *Eurylophella* | 0.25 | 0.80 | 0.32 | 0.75 |
| *Hydropsyche* | -1.07 | 0.44 | -2.39 | 0.02 |
| *Isonychia* | 0.26 | 0.55 | 0.46 | 0.64 |
| *Stenonema (Maccaffertium)* | 0.07 | 0.28 | 0.26 | 0.79 |
| *Nigronia* | -0.12 | 0.72 | -0.16 | 0.87 |
| *Psilotreta* | -2.76 | 0.94 | -2.91 | 0.003 |
| *Rhyacophila* | -0.25 | 1.00 | -0.25 | 0.80 |
| *Tipula* | -0.41 | 0.38 | -1.08 | 0.27 |
| *Tropisternus* | -1.06 | 0.68 | -1.57 | 0.11 |

Table S18: Results of Taxon-specific negative binomial GLMs testing the effect of restored sites on macroinvertebrate abundance in Spring 2025. Negative estimates indicate lower abundance post-translocation, while positive estimates represent higher abundance post-translocation. Models were fit separately for each taxon using a negative binomial error distribution to account for overdispersion.

| **Taxon** | **Estimate** | **SE** | **z (statistic)** | **p-value** |
| --- | --- | --- | --- | --- |
| *Amphinemura* | -3.64 | 1.05 | -3.46 | 0.0005 |
| *Anchytarsus* | -1.015 | 0.78 | -1.29 | 0.19 |
| *Calopteryx* | 7.73 | 0.61 | 1.27 | 0.20 |
| *Cheumatopsyche* | -0.54 | 0.40 | 1.34 | 0.18 |
| *Chimarra* | -2.48 | 1.04 | -2.39 | 0.02 |
| *Diplectrona* | -1.85 | 0.65 | -2.84 | 0.004 |
| *Ephemerella* | -1.50 | 0.45 | -3.28 | 0.001 |
| *Helichus* | -1.38 | 1.11 | -1.23 | 0.21 |
| *Hydropsyche* | -1.29 | 0.78 | -1.66 | 0.09 |
| *Ironoquia* | 2.76E-15 | 0.64 | 4.2868E-15 | 1 |
| *Isonychia* | -1.16 | 0.68 | -1.69 | 0.09 |
| *Stenonema (Maccaffertium)* | 0.78 | 0.51 | 1.51 | 0.13 |
| *Neophylax* | -3.55 | 1.04 | -3.40 | 0.0006 |
| *Nigronia* | -0.18 | 0.67 | -0.27 | 0.78 |
| *Penelomax* | -0.81 | 0.73 | -1.09 | 0.27 |
| *Plectrocnemia* | -0.39 | 0.35 | -1.11 | 0.26 |
| *Polycentropus* | 0.02 | 0.37 | 0.06 | 0.95 |
| *Psilotreta* | -1.83 | 0.84 | -2.18 | 0.02 |
| *Simulium* | -1.60 | 1.14 | -1.40 | 0.16 |
| *Timpanoga* | -0.34 | 0.45 | -0.75 | 0.44 |
| *Tipula* | 0.60 | 0.42 | 1.42 | 0.15 |

Table S19: PERMANOVA tests evaluating the effects of substrate type on macroinvertebrate community composition in Fall 2024 across all donor sites.

| **Source** | **df** | **Sum of Squares** | **R^2^** | **F** | **p-value** |
| --- | --- | --- | --- | --- | --- |
| Substrate type | 1 | 0.20198 | 0.06471 | 1.9371 | 0.107 |
| Residuals | 28 | 2.91954 | 0.93529 |  |  |
| Total | 29 | 3.12152 | 1.00 |  |  |

Table S20: PERMANOVA tests evaluating the effects of substrate type on macroinvertebrate community composition in Spring 2025 across all donor sites.

| **Source** | **df** | **Sum of Squares** | **R^2^** | **F** | **p-value** |
| --- | --- | --- | --- | --- | --- |
| Substrate type | 1 | 0.15728 | 0.05298 | 1.5663 | 0.144 |
| Residuals | 28 | 2.81161 | 0.94702 |  |  |
| Total | 29 | 2.96889 | 1.00 |  |  |

Table S21: PERMDISP results assessing homogeneity of multivariate dispersion between leaf and rock macroinvertebrate communities in Fall 2024 across donor sites

| Source | df | Sum of Squares | Mean Square | F | p-value |
| --- | --- | --- | --- | --- | --- |
| Site Type | 1 | 0.01667 | 0.01666 | 1.2052 | 0.2816 |
| Residuals | 28 | 0.38720 | 0.0138 |  |  |

Table S22: PERMDISP results assessing homogeneity of multivariate dispersion between leaf and rock macroinvertebrate communities in Spring 2025 across donor sites.

| Source | df | Sum of Squares | Mean Square | F | p-value |
| --- | --- | --- | --- | --- | --- |
| Site Type | 1 | 0.014969 | 0.014969 | 1.9242 | 0.1763 |
| Residuals | 28 | 0.217825 | 0.0077795 |  |  |

Table S23: Summary of Linear Models (LM) and a GLM testing the effect of substrate type (rock vs leaf) on different macroinvertebrate community metrics. Negative estimates indicate higher values on leaf substrate and vice versa

| Variable | Model Type | Estimate | SE | Test Statistic | df | p-value |
| --- | --- | --- | --- | --- | --- | --- |
| Taxa Richness | LM | -1.03 | 0.673 | -1.535 | 1 | 0.13 |
| Shannon Diversity | LM | 0.173 | 0.119 | 1.455 | 1 | 0.151 |
| EPT Richness | GLM (Poisson) | 0.04 | 0.10 | 0.407 |  | 0.684 |
| Abundance | LM | -45 | 19.99 | -2.251 | 1 | 0.02 |
